# PhenoPlasm: a database of disruption phenotypes for malaria parasite genes

**DOI:** 10.1101/101717

**Authors:** Theo Sanderson, Julian C. Rayner

**Affiliations:** Wellcome Trust Sanger Institute, Hinxton, Cambridge

## Abstract

Two decades from the first *Plasmodium* transfection, attempts have been made to disrupt more than 900 genes in malaria parasites, across five *Plasmodium* species. While results from the rodent malarias have been curated and systematised, phenotypic data for species of human malaria parasites has existed only scattered across a large literature. To facilitate systematic views of known experimental-genetic data across *Plasmodium* species, we have built PhenoPlasm (<u>http://www.phenoplasm.org</u>), a database of phenotypes for *Plasmodium* parasites. The site provides a simple interface to link citation-backed *Plasmodium* reverse-genetic phenotypes to gene IDs. The database has been populated with phenotypic data on 330 *P. falciparum* genes, curated from 155 individual publications, as well as existing curated data from RMgmDB. These data are presented using 1: 1 ortholog mapping to allow a researcher interested in a gene in one species to see results across *Plasmodium.* The collaborative nature of the database enables any researcher to add new phenotypes as they are discovered.

## Introduction

The increasing use of reverse genetics in *Plasmodium spp.* has provided countless insights into the biology of the malaria parasite (de Koning-Ward, Gilson, & Crabb, 2015). Nevertheless, to date such results are (with the notable exception of rodent malaria species) scattered across a vast literature. This means that a researcher whose experiment or analysis reveals a set of genes must devote considerable time to reviewing papers if they are to understand what is known about their knock-out phenotypes. To facilitate rapid functional profiling using already established phenotypes, we set out to build a database to contain this information.

There were three key functional requirements for such a database:

### Systematic, and synergistic with existing resources

To allow automated bioinformatic analyses, it is crucial that the database have a defined, machine-comprehensible, schema for recording phenotypes. It is also important that this schema is compatible with existing resources.

The rodent malaria genetically modified parasite database (RMgmDB, <u>http://pberghei.eu</u>; Khan, Kroeze, Franke-Fayard, & Janse, 2013) provides a powerful curated resource for the rodent malarias. Nevertheless, there is a need to bring its results together with those from human-infecting systems to provide a comprehensive view of the genes likely to be required for different aspects of parasite biology in the human malaria parasites, some of which lack rodent orthologs. To allow such integration, any new database schema must be compatible with that of RMGMdb, which comprises 6 different stages at which phenotypes can occur (asexual, gametocyte/gamete, fertilization & ookinete, oocyst, sporozoite and liver). RMgmDB also distinguishes cases in which a modification is not successful, which provide some implication of a possible role in asexual growth; we decided to call this quality *mutant viability,* though of course the failure to obtain a mutant might also result from a technical failure.

### Orthology-based retrieval

The use of model systems has strongly facilitated attempts to understand human malaria parasites (Zuzarte-Luis, Mota, & Vigario, 2014). These systems are valuable both because rodent malaria parasites appear to be technically simpler to genetically modify and because their *in vivo* nature has allowed the study of some aspects of parasite biology in a more physiological setting. Rodent models make the recapitulation of the complete lifecycle much more achievable and so comprise the vast majority of known non-blood-stage phenotypes. Nevertheless, murine parasites do not contain orthologs of every *P. falciparum* gene and so these studies cannot provide a complete view of the genetics of the parasites causing human disease, and even where orthologs exist their phenotypes may not always be conserved.

We decided that an optimal approach would allow a researcher to search for gene IDs from any species but at a glance to see both results in this species and for orthologous genes in other *Plasmodium* species. The database should contain records for emerging new genetic models, such as *P. knowlesi* (Kocken et al., 2002; Moon et al., 2013), as well as *P. vivax,* which though currently genetically intractable is of key medical importance and where gene functions may be interpreted through orthology.

### Crowd-sourced, or with the potential to be

There are a number examples of previous databases which have become out of date as the enthusiasm, or funding, for curating them has ceased. Creating a database which allows user submissions theoretically allows the database to be maintained primarily by the research-community. This requires the creation of an intuitive interface for entry of data and means that the primary data source is publications, with each phenotype backed up with a PubMed ID.

## Methods

### Database schema

The database comprises 3 main data tables.

### Genes

This table contains data genes IDs, the species to which they belong, their gene symbol, product description and ortholog group (created by EuPathDB using OrthoMCL). This data comes from PlasmoDB (and the gene models originally from GeneDB). (Aurrecoechea et al., 2009; Chen, Mackey, Stoeckert, & Roos, 2006; Logan-Klumpler et al., 2012)

### Aliases

This table contains a record for each possible identity that could be used to refer to gene (i.e. historical IDs) and links these to the gene record.

### Phenotypes

This table contains a record for every described phenotype. Each record corresponds to a stage of the lifecycle and provides a phenotype from a predefined list (e.g. *no difference, attenuated, egress defect),* backed up by a citation. The type of experimental approach used to disrupt the gene is also recorded, allowing a distinction between conditional methods, which provide strong evidence of essentiality, from a failure to modify the locus, which provides weaker evidence.

### Display

The database can be queried either for a set of genes or a single gene. The former provides a table with one line per gene while the latter provides referencing for each claim and displays any additional notes.

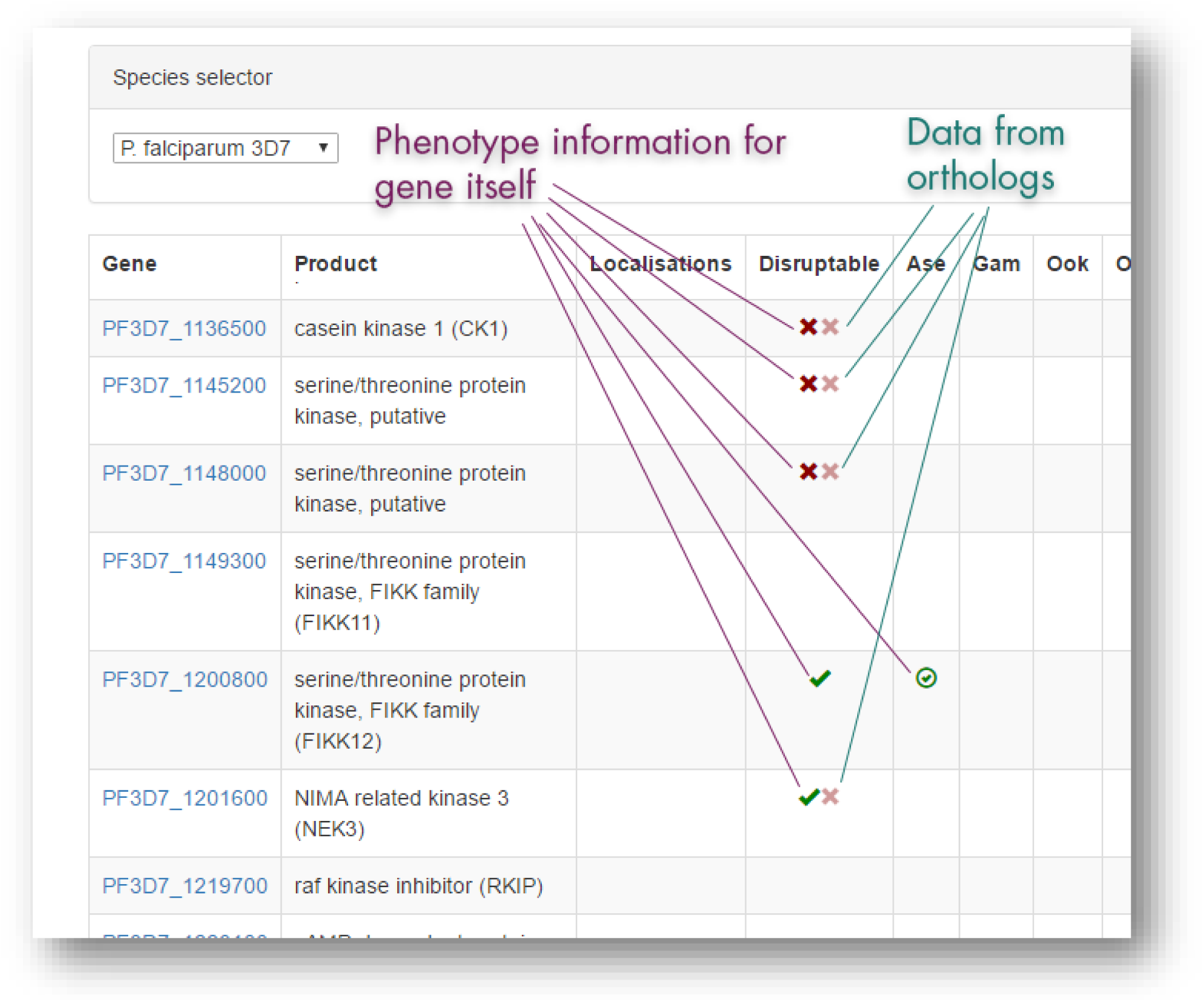

### Literature review

A scan was made of the *Plasmodium* literature using Google Scholar (which provides full-text search for a large proportion of publications). Terms for which complete literature scans were made included ‘“attempts to disrupt” falciparum’, ‘“gene disruption of” falciparum’ ‘“gene deletion construct” falciparum’ and ‘“gene disruption construct” falciparum’. A number of other terms were used, but with results only curated for the first 10 pages of results. In addition, genes with a suggested role in erythrocyte invasion were systematically curated by searching for all references to any version of their gene IDs, as discussed below.

One challenge in conducting literature searches into *Plasmodium* proteins is the fact that the numerous iterative improvements made to *Plasmodium* genome assemblies mean that a gene could be referred to by any of numerous historic identifiers. To assist with this, the PhenoPlasm page for each gene interface contains a button which conducts a boolean search on Google Scholar, searching article full text for *any* of the identifiers which have been used to refer to the gene.

All genes were initially been marked as ‘unresearched’ to warn the end-user against trusting the resource to be fully-exhaustive. As some genes have been researched by a thorough literature review, this warning has been removed from them.

## Results

Some form of phenotyping information is available for 909 genes, 576 are from rodent malaria parasites and so represent data imported from RMGMdb, leaving 333 non-rodent parasite genes which are the result of our curation.

There are 3708 total phenotyping datapoints (one life-stage, for one gene knock-out, in one study). A large majority of these are from the rodent parasites, since relatively few P. falciparum genes have phenotyping data reported beyond the blood stage.

Since there is overlap between the species, the number of ortholog groups covered in at least one species is 672. This represents approximately 15% of the core *Plasmodium* genome.

| Organism | Number of genes w/ phenotyping data | Main source |
| --- | --- | --- |
| *P. falciparum* | 330 | Literature curation |
| *P. knowlesi* | 3 | Literature curation |
| *P. berghei* | 523 | RMgmDB |
| *P. yoelii* | 53 | RMgmDB |
| Total | 909 | All |

## Discussion

*Plasmodium* reverse genetics is now beginning to reach the genome-level stage. As it does so, phenotypic data has the potential to shed light on the importance of the novel genes found in these early-branching eukaryotes.

Large scale genetic modification programmes in *Plasmodium* are already underway (Bronner et al., 2016; Gomes et al., 2015) and so the 85% of the genome for which there are no published phenotypes should be a portion that shrinks rapidly. Nevertheless, no single approach is likely to reach saturation for some time, and exploring the complete parasite lifecycle is likely to take much longer. In addition, the lack of non-homologous end-joining pathway in *Plasmodium* parasites limits the use of the conventional CRISPR-Cas9 screens, which have revolutionized genetics in other organisms. For these reasons, a complete view of the reverse-genetic landscape for a gene or pathway will require bringing together multiple datasets with the individual gene-by-gene studies that have characterized decades of research.

We hope that PhenoPlasm assists in these developments and that it allows the prioritization of future large scale studies, by eliminating duplication of existing efforts, and allowing a focus on the portion of the genome which remains wholly unexplored by reverse genetic approaches. These genes of unknown function may yet contain crucial for parasite virulence and perhaps druggable targets.

